# Mutations in *TAC1B*: a novel genetic determinant of clinical fluconazole resistance in *C. auris*

**DOI:** 10.1101/2020.02.18.955534

**Authors:** Jeffrey M. Rybak, José F. Muñoz, Katherine S. Barker, Josie E. Parker, Brooke D. Esquivel, Elizabeth L. Berkow, Shawn R. Lockhart, Lalitha Gade, Glen E. Palmer, Theodore C. White, Steve L. Kelly, Christina A. Cuomo, P. David Rogers

## Abstract

*Candida auris* has emerged as a multidrug-resistant pathogen of great clinical concern. Approximately 90% of clinical *C. auris* isolates are resistant to fluconazole, the most commonly prescribed antifungal agent, yet it remains unknown what mechanisms underpin this fluconazole resistance. To identify novel mechanisms contributing to fluconazole resistance in *C. auris*, the fluconazole-susceptible *C. auris* clinical isolate AR0387 was passaged in media supplemented with fluconazole to generate derivative strains which had acquired increased fluconazole resistance *in vitro*. Comparative analysis of comprehensive sterol profiles, [^3^H]-fluconazole uptake, sequencing of *C. auris* genes homologous to genes known to contribute to fluconazole resistance in other species of *Candida*, and the relative expression of *C. auris ERG11, CDR1*, and *MDR1* were performed. All fluconazole-evolved derivative strains were found to have acquired mutations in the zinc-cluster transcription factor-encoding gene, *TAC1B*, and a corresponding increase in *CDR1* expression relative to the parental clinical isolate, AR0387. Mutations in *TAC1B* were also identified in a set of 304 globally distributed *C. auris* clinical isolates representing each of the four major clades. Introduction of the most common mutation found among fluconazole-resistant clinical isolates of *C. auris* into the fluconazole-susceptible isolate AR0387, was confirmed to increase fluconazole resistance by 8-fold, and the correction of the same mutation in a fluconazole-resistant isolate, AR0390, decreased fluconazole MIC by 16-fold. Taken together, these data demonstrate that *C. auris* can rapidly acquire resistance to fluconazole *in-vitro*, and that mutations in *TAC1B* significantly contribute to clinical fluconazole resistance.

**IMPORTANCE:** *Candida auris* is an emerging multidrug-resistant pathogen of global concern, known to be responsible for outbreaks on six continents and commonly resistant to antifungals. While the vast majority of clinical *C. auris* isolates are highly resistant to fluconazole, an essential part of the available antifungal arsenal, very little is known about the mechanisms contributing to resistance. In this work, we show that mutations in the transcription factor *TAC1B* significantly contribute to clinical fluconazole resistance. These studies demonstrate that mutations in *TAC1B* can arise rapidly in vitro upon exposure to fluconazole, and that a multitude of resistance-associated *TAC1B* mutations are present among the majority of fluconazole-resistant *C. auris* isolates from a global collection and appear specific to a subset of lineages or clades. Thus, identification of this novel genetic determinant of resistance significantly adds to the understanding of clinical antifungal resistance in *C. auris*.

## INTRODUCTION

First identified in 2009, *Candida auris* has rapidly become a healthcare-associated and multidrug-resistant pathogen of global concern.(1, 2) While originally found to be the causative pathogen of virtually simultaneous outbreaks of invasive candidiasis in Asia, South Africa, and South America, *C. auris* has now been identified in more than 30 countries across 6 continents, including more than 900 confirmed clinical cases of *C. auris* infections in the United States.(3) Further contributing to the clinical significance of this organism are its proclivity to colonize both environmental surfaces and patients, challenges associated with the reliable identification in the clinical microbiology laboratory, and the markedly decreased susceptibility to currently available antifungal agents found among a large proportion of *C. auris* clinical isolates.(4, 5) While epidemiologic data and clinical experience pertaining to the treatment of infections caused by *C. auris* are currently inadequate to support the establishment of epidemiologic cut-off values and true clinical breakpoints, the Center for Disease Control and Prevention (CDC) has proposed tentative breakpoints to help guide clinicians based upon available susceptibility data for *C. auris* clinical isolates. When these tentative breakpoints are applied, approximately 3% of *C. auris* clinical isolates are resistant to echinocandins, one third are resistant to amphotericin B, and 90% are resistant to fluconazole (minimum inhibitory concentration [MIC] ≥32mg/L; modal MIC ≥256mg/L).(6) Additionally, one third of clinical isolates are multidrug-resistant, with elevated MIC for two or more different classes of antifungals, and clinical isolates resistant to all available agents have been repeatedly reported.(7, 8)

The extent of fluconazole resistance among *C. auris* isolates is particularly concerning as this agent remains the most commonly prescribed antifungal, and many of the outbreaks of *C. auris* have occurred in resource-limited settings.(2, 8-11) While the pervasiveness of fluconazole resistance among *C. auris* clinical isolates substantially limits therapeutic options of *C. auris* infections, relatively little is known about the molecular mechanisms underpinning this resistance. One mechanism of fluconazole resistance repeatedly identified in *C. auris* is mutation of the gene encoding the sterol-demethylase enzyme targeted by the triazoles, *ERG11*. Three such mutations, encoding the amino acid substitutions VF125AL (commonly referred to as F126L), Y132F, and K143R, are frequently reported among fluconazole-resistant clinical isolates, and associations between these mutations and specific genetic clades of *C. auris* have been observed.(2) Additionally, the mutations encoding the Y132F and K143R substitutions correspond to mutations known to contribute to triazole resistance in other species of *Candida* such as *Candida albicans*.(12) While the direct impact of these *ERG11* mutations has not been delineated in *C. auris*, heterologous expression of *C. auris ERG11* alleles carrying mutations encoding either the Y132F or K143R amino acid substitutions on a low copy number episomal plasmid was observed to decrease fluconazole susceptibility in a haploid strain of *Saccharomyces cerevisiae*.(13) However, clinical isolates harboring the same *ERG11* mutations and exhibiting fluconazole MIC as low as 1mg/L have been described, as have fluconazole-resistant isolates of *C. auris* with no mutation in *ERG11*, suggesting the presence of yet to be identified mechanisms of fluconazole resistance.(8, 14)

In addition to mutations in *ERG11*, increased expression of efflux pump-encoding genes is a common contributor to clinical triazole resistance among multiple species of *Candida*.(15) Most notable of these is *C. glabrata*, in which nearly all of clinical triazole resistance is attributable to overexpression of the ATP-Binding Cassette (ABC)-type efflux pump encoding genes *CgCDR1, CgPDH1*, and *CgSNQ2*.(16) The *C. auris* genome has recently been revealed to encode a substantial number of efflux pump encoding genes of both the ABC and Major Facilitator Superfamily (MFS) classes, and triazole-resistant isolates of *C. auris* have been observed to exhibit efflux pump activity greatly exceeding (up to 14-fold higher) that of *C. glabrata*.(17-19) Furthermore, the increased expression of the *C. auris* ABC-type efflux pump-encoding gene, *CDR1*, has previously been shown to substantially contribute to clinical triazole resistance.(20, 21) At present however, the genetic determinants underpinning the increased expression of efflux pump-encoding genes in *C. auris* remain unidentified.

In this work, we undertook an unbiased approach utilizing *in-vitro* evolution to create a collection of isogenic *C. auris* strains with increased fluconazole resistance, exhibiting an 8 to 64-fold increase in fluconazole MIC. Characterization of these strains as well as analysis of whole genome sequencing data for over 300 globally-distributed *C. auris* isolates implicated *TAC1B* (B9J08_004820), a close homolog of the well-characterized *C. albicans* transcriptional regulator *CaTAC1*, as a novel genetic determinant of clinical fluconazole resistance. Having identified *TAC1B* mutations to be present among a large proportion of fluconazole-resistant clinical isolates, we utilized a Cas9-mediated transformation system to both introduce the most common *TAC1B* mutation identified among resistant clinical isolates (encoding A640V) into the fluconazole-susceptible AR0387, as well as correct the A640V-encoding mutation in the previously characterized and highly fluconazole-resistant clinical isolate AR0390 to the wild-type sequence. In both cases, the presence of this prevalent *TAC1B* mutation was associated with significant increase in fluconazole MIC, demonstrating that mutations in *TAC1B* represent a prevalent and significant genetic determinant of fluconazole resistance among clinical *C. auris* isolates.

## RESULTS

### *Candida auris* rapidly acquires increased fluconazole resistance *in-vitro*

In an effort to identify novel mechanisms of fluconazole resistance in this emerging multidrug-resistant pathogen, a collection of isogenic strains with increased fluconazole resistance was created via *in-vitro* evolution utilizing the previously described fluconazole-susceptible *C. auris* clinical isolate AR0387 (also known as B8441) (**Figure 1**). Briefly, the parental AR0387 was grown in liquid cultures of YPD media supplemented with either 8 or 32 mg/L of fluconazole for 48 hours. Each liquid culture was then plated on the standard antifungal susceptibility testing media, RPMI, supplemented with the same concentration of fluconazole for an additional 48 hours to identify individual colonies exhibiting increased fluconazole resistance. Two individual colonies were selected for characterization from the plate supplemented with 8mg/L of fluconazole (yielding strains FLU-A and FLU-B), and a single colony was selected from the plate supplemented with 32mg/L of fluconazole (yielding strain FLU-C). One strain from each initial passage, FLU-A and FLU-C, were subsequently subjected to a second passage in 64 and 256mg/L of fluconazole supplemented media, respectively, yielding strains FLU-A2 and FLU-C2.

**Figure 1.**
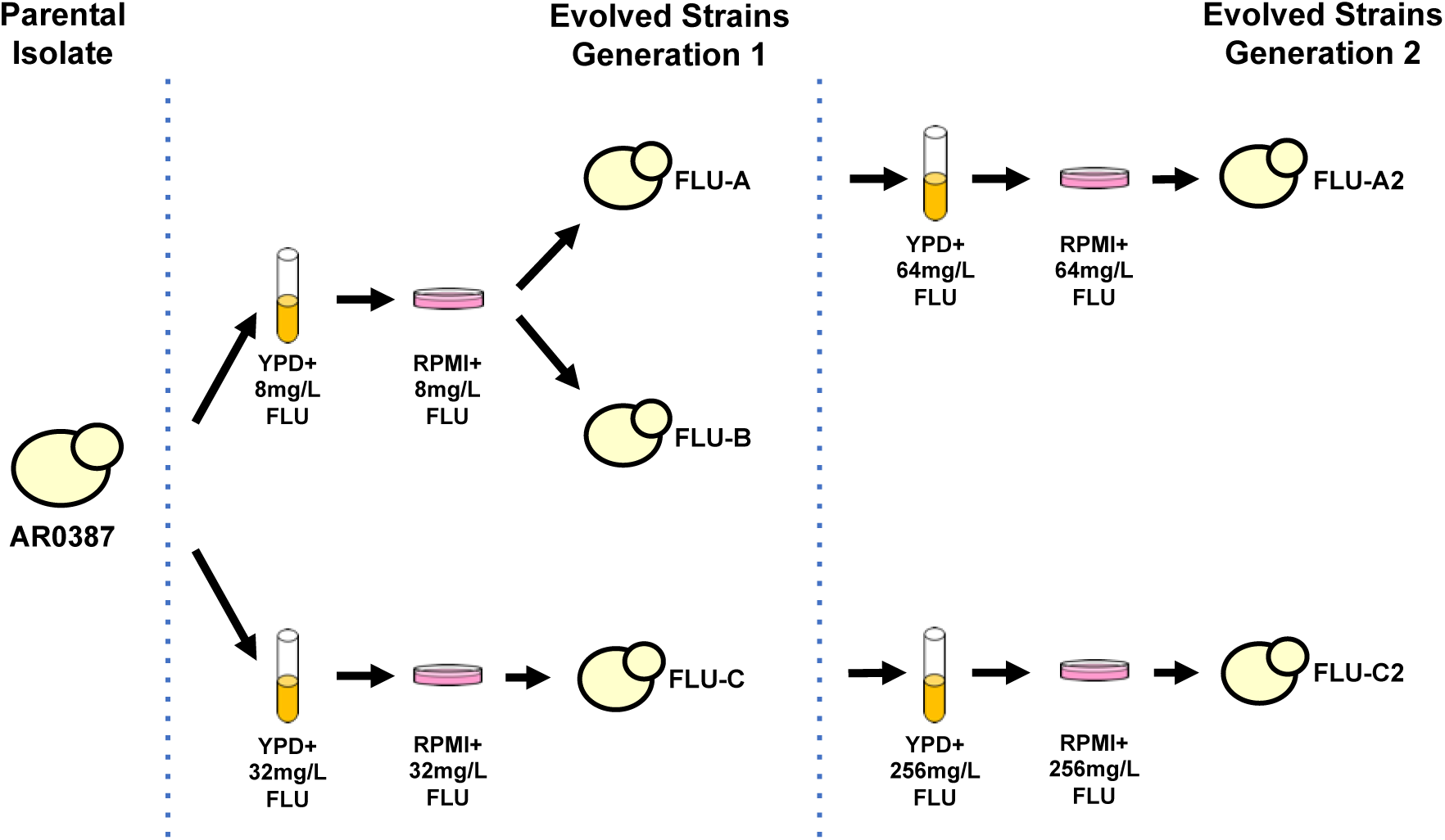
Schematic of *C. auris* fluconazole *in-vitro* evolution experiments. AR0387 was cultured in YPD supplemented with 8 or 32mg/L of fluconazole. Cultures were plated on RPMI containing the same concentration of fluconazole, and individual colonies were picked for further characterization. Fluconazole-evolved strains FLU-A and FLU-C were subsequently further passaged in YPD supplemented with 64 and 256mg/L of fluconazole, respectively. Cultures were then again plated on RPMI containing the same concentration of fluconazole and individual colonies were picked for further characterization.

Fluconazole MIC were then determined for the parental AR0387 and each of the five fluconazole-evolved strains by broth microdilution in accordance with Clinical Laboratory Standards Institute methodology with minor modifications as previously described.(20) AR0387 exhibited a fluconazole MIC of 1mg/L, while the five fluconazole-evolved strains were found to have MIC ranging from 8 to 64mg/L (**Figure 2**). Each of the second-generation evolved strains, FLU-A2 and FLU-C2, exhibited a further 2 to 4-fold increase in fluconazole MIC relative to their respective first-generation strains.

**Figure 2.**
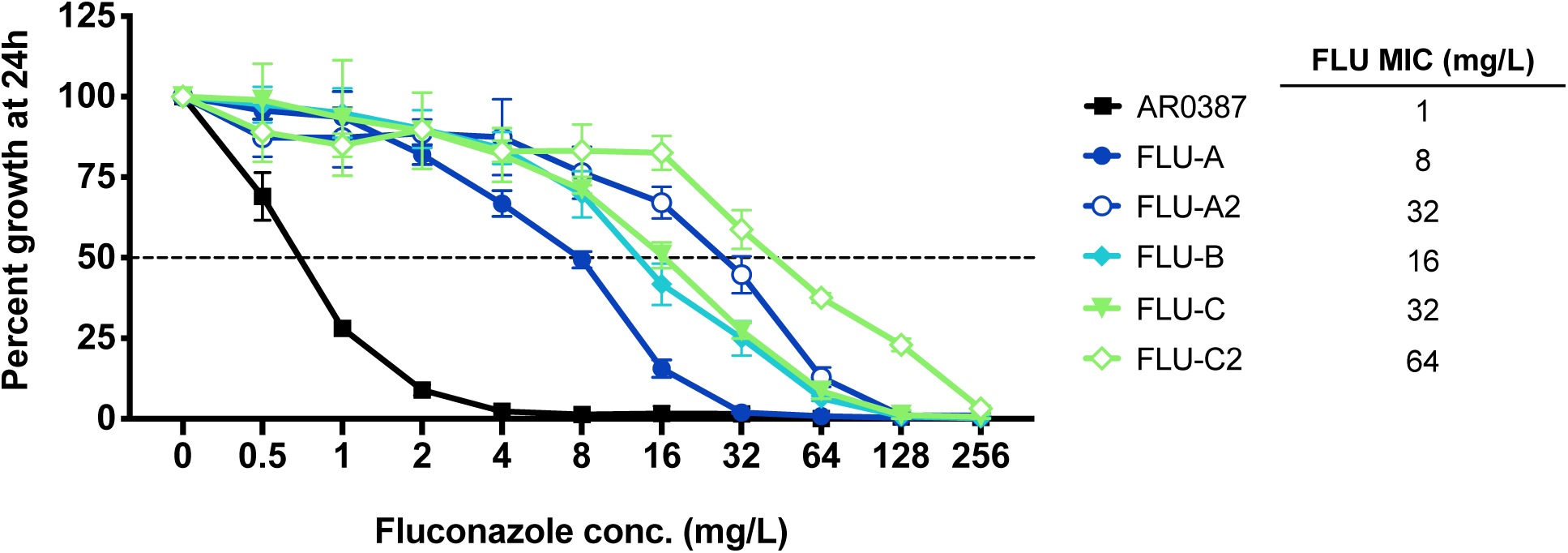
MIC of *C. auris* fluconazole-evolved strains. Percentage growth of AR0387 and fluconazole-evolved strains with escalating concentrations of fluconazole measured at 24 hours. Percent growth was determined relative to respective untreated controls as assessed by absorbance at OD_600_.

### Fluconazole-evolved strains exhibit alterations in membrane sterols without accompanying mutations in *ERG11* or *ERG3*

As fluconazole-resistant *C. auris* clinical isolates are very often found to possess mutations in *ERG11*, sequencing of the *ERG11* allele for each of the fluconazole-evolved strains was performed. Surprisingly, all evolved strains were found to have wildtype *ERG11* sequences matching that of the parental AR0387. To assess for other changes to the ergosterol biosynthesis pathway which may be contributing to fluconazole resistance, each of the fluconazole-evolved strains and the parental AR0387 were subsequently subjected to comprehensive sterol profiling. Briefly, each strain was grown to exponential growth phase in RPMI liquid media with or without 16mg/L of fluconazole (a concentration approximating the average serum concentration achieved in patients being treated for candidemia).(23)

Following growth in RPMI without fluconazole, all fluconazole-evolved strains and the parental AR0387 were observed to have largely similar sterol profiles (**Figure 3**). In all samples, ergosterol comprised more than 75% of total cellular sterols, with ergosta-5, 7, 22, 24(28)-tetraenol and zymosterol observed to be the next most abundant sterols. Intriguingly, four of the fluconazole-evolved strains (FLU-A, FLU-B, FLU-C, and FLU-A2) were also observed to have a small amount (2 to 4%) of 14-methyl-fecosterol present, while this sterol was absent in both AR0387 and FLU-C2. In *Candida albicans*, 14-methyl-fecosterol is a known substrate of the sterol-desaturase enzyme encoded by *CaERG3*, which catalyzes the conversion of 14-methyl-fecosterol to the toxic sterol associated with the antifungal activity of the triazoles, 14-methyl-ergosta-8, 24(28)-dienol-3,6-diol (**Figure 4**).

**Figure 3.**
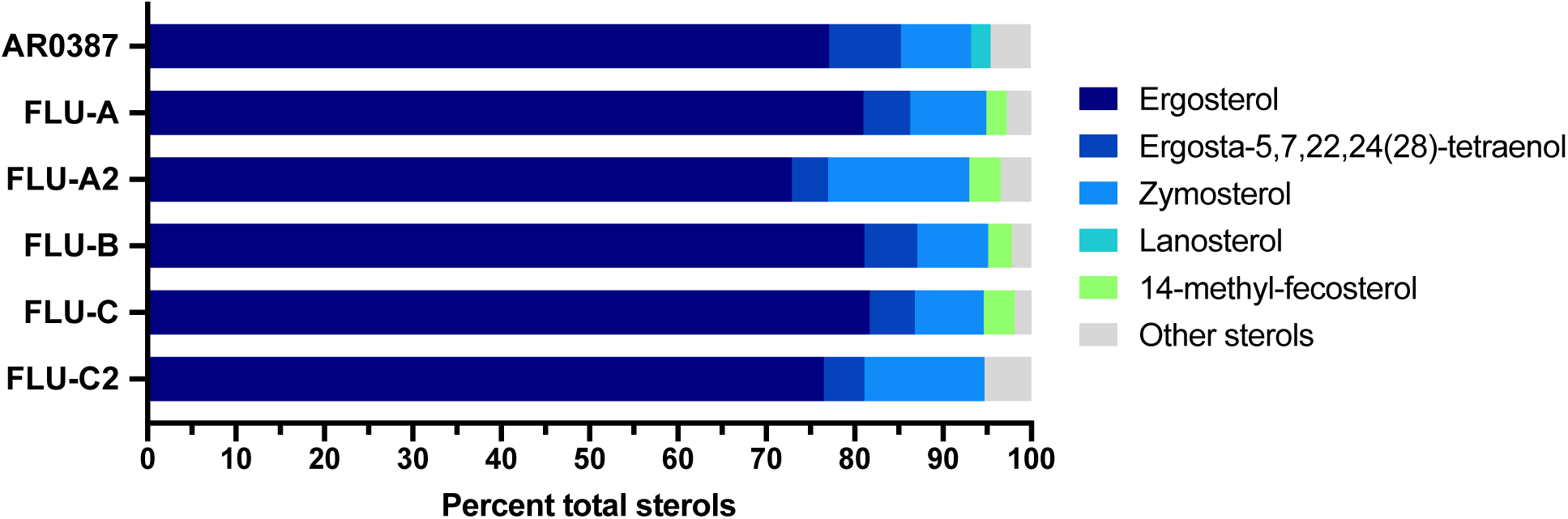
Sterol profiles of *C. auris* evolved strains. The major constituent sterols for AR0387 and fluconazole-evolved strains at exponential growth phase in RPMI are shown as the proportion of total cellular sterols.

**Figure 4.**
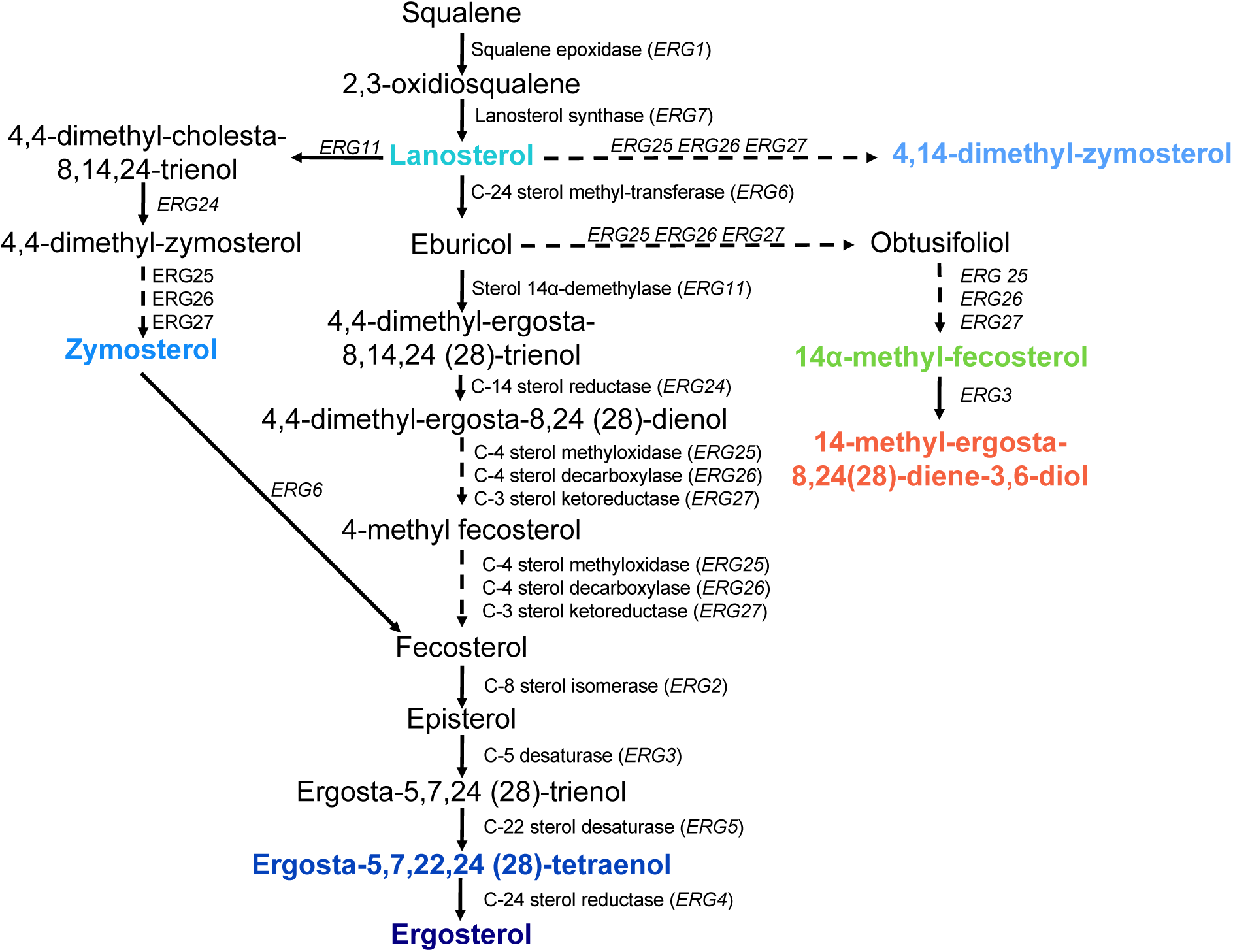
Predicted *C. auris* sterol biosynthesis pathway. The major constituent sterols identified in sterol profiles are shown with corresponding colors.

Following growth in RPMI supplemented with fluconazole, the sterol profiles of each of the fluconazole-evolved strains were dramatically different than that of AR0387 (**Figure 5**). While ergosterol was still the predominant sterol among all five fluconazole-evolved strains, lanosterol (46%) and 14-methyl-fecosterol (21%) were observed to be the two most prevalent sterols in AR0387. Additionally, 14-methyl-ergosta-8, 24(28)-dienol-3,6-diolcomprised 4% all sterols present in AR0387, while this sterol was absent in the sterol profiles of all fluconazole-evolved strains. As mutations in the sterol-desaturase encoding gene, *ERG3*, have been observed to contribute to fluconazole resistance in other species of *Candida*, and notable differences in the amount of cellular 14-methyl-fecosterol and 14-methyl-ergosta-8, 24(28)-dienol-3,6-diol were observed between AR0387 and the fluconazole-evolved strains, sequencing of the *C. auris* gene (B9J08_003737) with the highest degree of homology to *C. albicans CaERG3* was performed. However, no mutation in *C. auris ERG3* was observed in any of the fluconazole-evolved strains.

**Figure 5.**
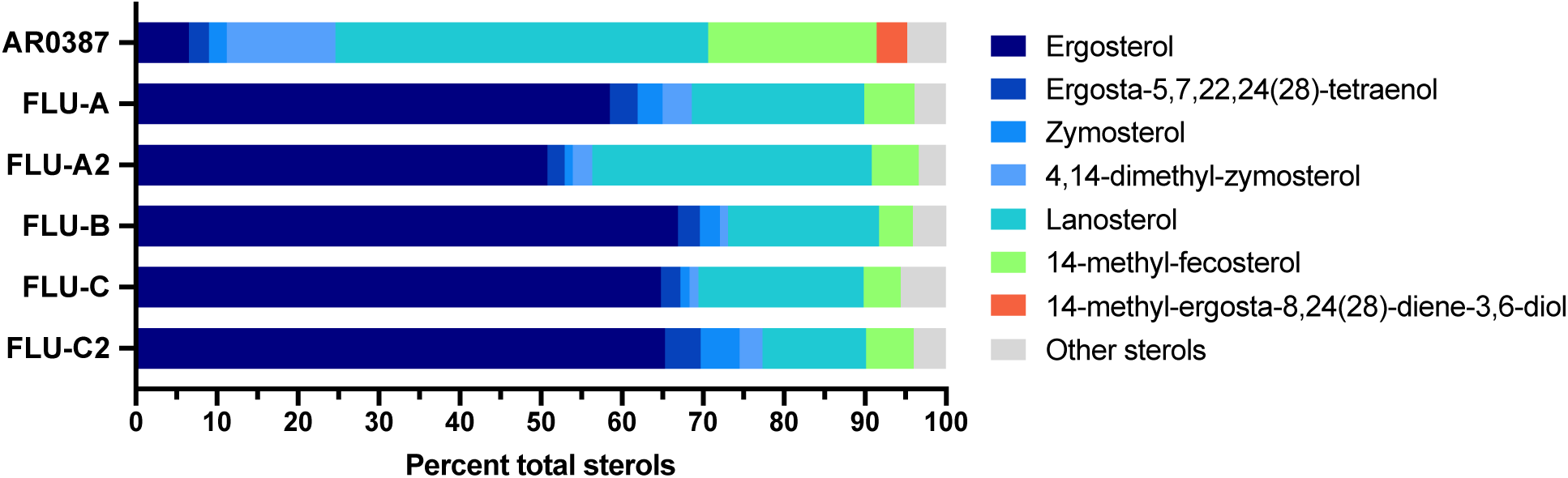
Sterol profiles of *C. auris* evolved strains with fluconazole exposure. The major constituent sterols for AR0387 and fluconazole-evolved strains at exponential growth phase in RPMI supplemented with 16mg/L fluconazole are shown as the proportion of total cellular sterols.

### Fluconazole-evolved strains exhibit significantly reduced fluconazole uptake

As triazoles including fluconazole have previously been shown to enter the cells of *C. albicans* via facilitated diffusion, deficient drug import was next examined as a potential mechanism contributing to the increased fluconazole resistance among the fluconazole-evolved strains.(24) As previously described, the accumulation of [^3^H]-fluconazole was assessed for AR0387 and each fluconazole-evolved strain, as well as a previously characterized strain of AR0387 where the *CDR1* gene has been deleted (AR0387_Δ*cdr1*), following a two hour glucose starvation in YNB media without carbon source supplementation.(20) [^3^H]-fluconazole accumulation was observed to be reduced by approximately 50% in four of the fluconazole-evolved strains (FLU-A, FLU-B, FLU-C, and FLU-A2) relative to than that observed in AR0387 (**Figure 6A**), while accumulation in FLU-C2 did not significantly differ from that of AR0387. Importantly, there was no difference in [^3^H]-fluconazole accumulation between AR0387 and AR0387_Δ*cdr1* (**Figure 6B**), confirming that the conditions used in this study for glucose starvation were adequate to remove the activity of this known *C. auris* resistance effector.

**Figure 6.**
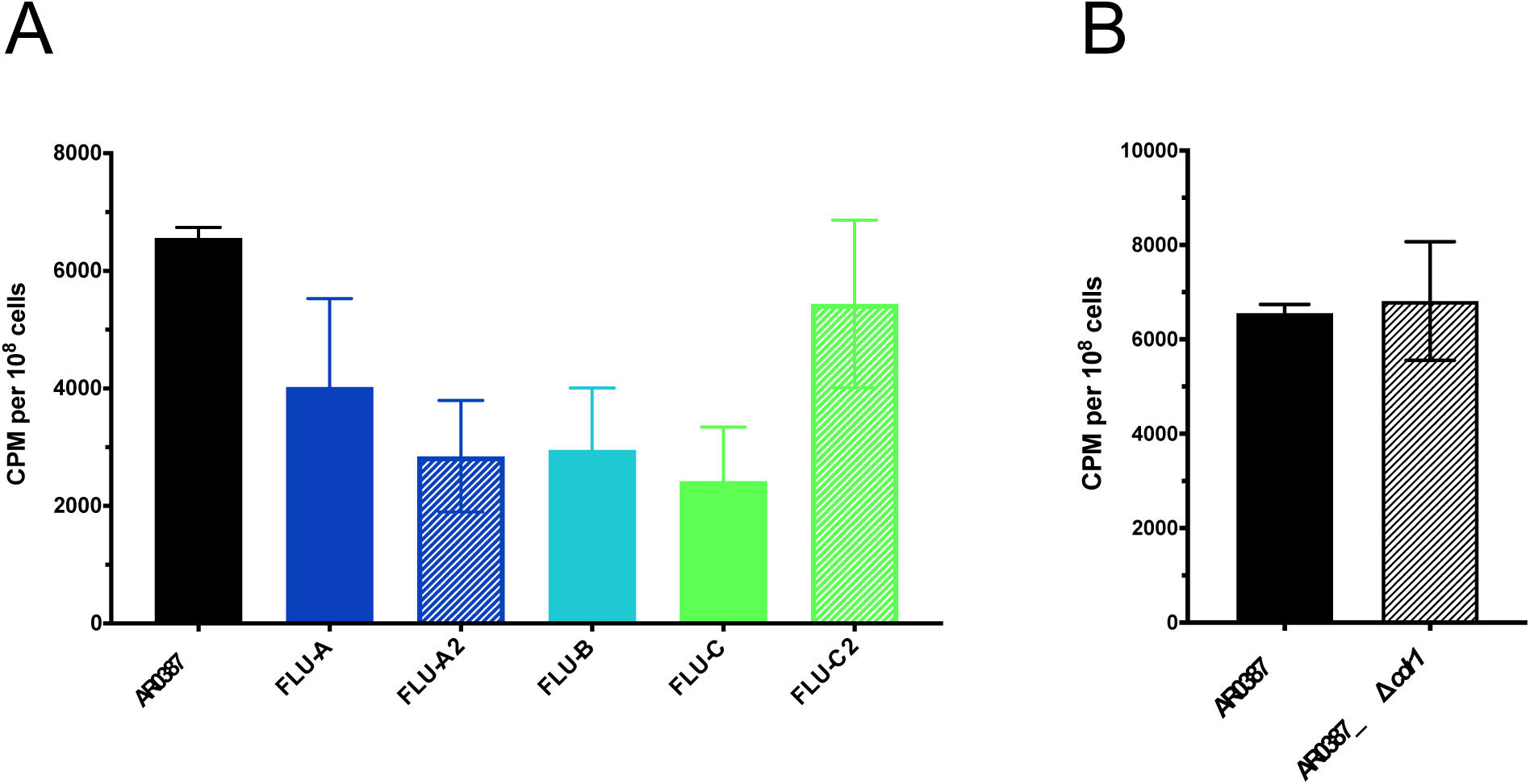
Intracellular accumulation of [^3^H]-fluconazole among fluconazole-evolved strains. [^3^H]-labeled fluconazole uptake in (a) fluconazole-evolved strains and (b) *CDR1-*deletion strain, compared to the parental clinical isolate AR0387.

### Mutations in *TAC1B* are associated with significantly increased expression of *CDR1*

Gain-of-function mutations in zinc-cluster transcription factor genes, such as *C. albicans* genes *CaUPC2, CaMRR1*, and *CaTAC1* are a well-characterized mechanism of fluconazole resistance among other species of *Candida*.(15) To determine if similar mutations may be contributing to the fluconazole resistance among the fluconazole-evolved strains in these studies, the *C. auris* genes with the highest degree of homology to the *C. albicans* transcriptional regulatory genes *CaUPC2, CaMRR1*, and *CaTAC1*, here named *UPC2* (B9J08_000270), *MRR1* (B9J08_004061), *TAC1A* (B9J08_004819), and *TAC1B* (B9J08_004820), were identified by BLAST and gene orthology analysis and sequencing was performed. As two *C. auris* genes possessing very high degrees of homology with *CaTAC1* were identified, both were included in this study. While no mutations were identified in *TAC1A* or *MRR1*, all five fluconazole-evolved strains were found to have mutations encoding amino acid substitutions in *TAC1B* (**Table 1**). Both the FLU-A strain its second-generation derivative FLU-A2 had acquired a mutation encoding the amino acid substitution R495G, while the strains FLU-B, FLU-C, and FLU-C2 all had acquired a mutation encoding the amino acid substitution F214S. Neither of these mutations correspond to previously characterized GOF mutations in *CaTAC1*, or orthologous genes from other species of *Candida*. However, these mutations are predicted to alter residues near or within the conserved fungal transcription factor middle homology region (MHR) of Tac1Bp, and multiple mutations encoding amino acid substitutions in the MHR of *CaTAC1* have previously been associated with fluconazole resistance.(25) Additionally, a sole mutation in *UPC2* encoding the amino acid substitution M365I was identified in FLU-C2, and this mutation similarly alters a residue predicted to reside within the MHR of Upc2p.

**Table 1.** Sequencing of *C. auris MRR1, TAC1A, TAC1B*, and *UPC2* among fluconazole-evolved strains.

| Gene | AR0387 | FLU-A | FLU-A2 | FLU-B | FLU-C | FLU-C2 |
| --- | --- | --- | --- | --- | --- | --- |
| *MRR1* | WT | WT | WT | WT | WT | WT |
| *TAC1A* | WT | WT | WT | WT | WT | WT |
| *TAC1B* | WT | **R495G** | **R495G** | **F214S** | **F214S** | **F214S** |
| *UPC2* | WT | WT | WT | WT | WT | **M365I** |

In an effort to ascertain if the identified mutations in *TAC1B* and *UPC2* may be associated with altered expression of potential resistance effectors among the fluconazole-evolved strains, the relative expression of *ERG11, CDR1*, and *MDR1* was evaluated by RTqPCR. To accomplish this, AR0387 and each of the fluconazole-evolved strains were grown to exponential growth phase in RPMI media, and RNA was extracted as previously described. The expression of each gene of interest relative to AR0387 was assessed using the *ΔΔCT* (threshold cycle) method and the *C. auris ACT1* housekeeping gene (B9J08_000486).(20) Among all five fluconazole-evolved strains, the expression of *CDR1* was found to be 3 to 5-fold that of AR0387 (**Figure 7**). This level of *CDR1* expression is similar to that previously described among extensively fluconazole-resistant *C. auris* clinical isolates.(20) Additionally, subtle variations in the expression of *ERG11*, and *MDR1*, not exceeding 2.1-fold that of AR0387, were also observed among individual fluconazole-evolved strains.

**Figure 7.**
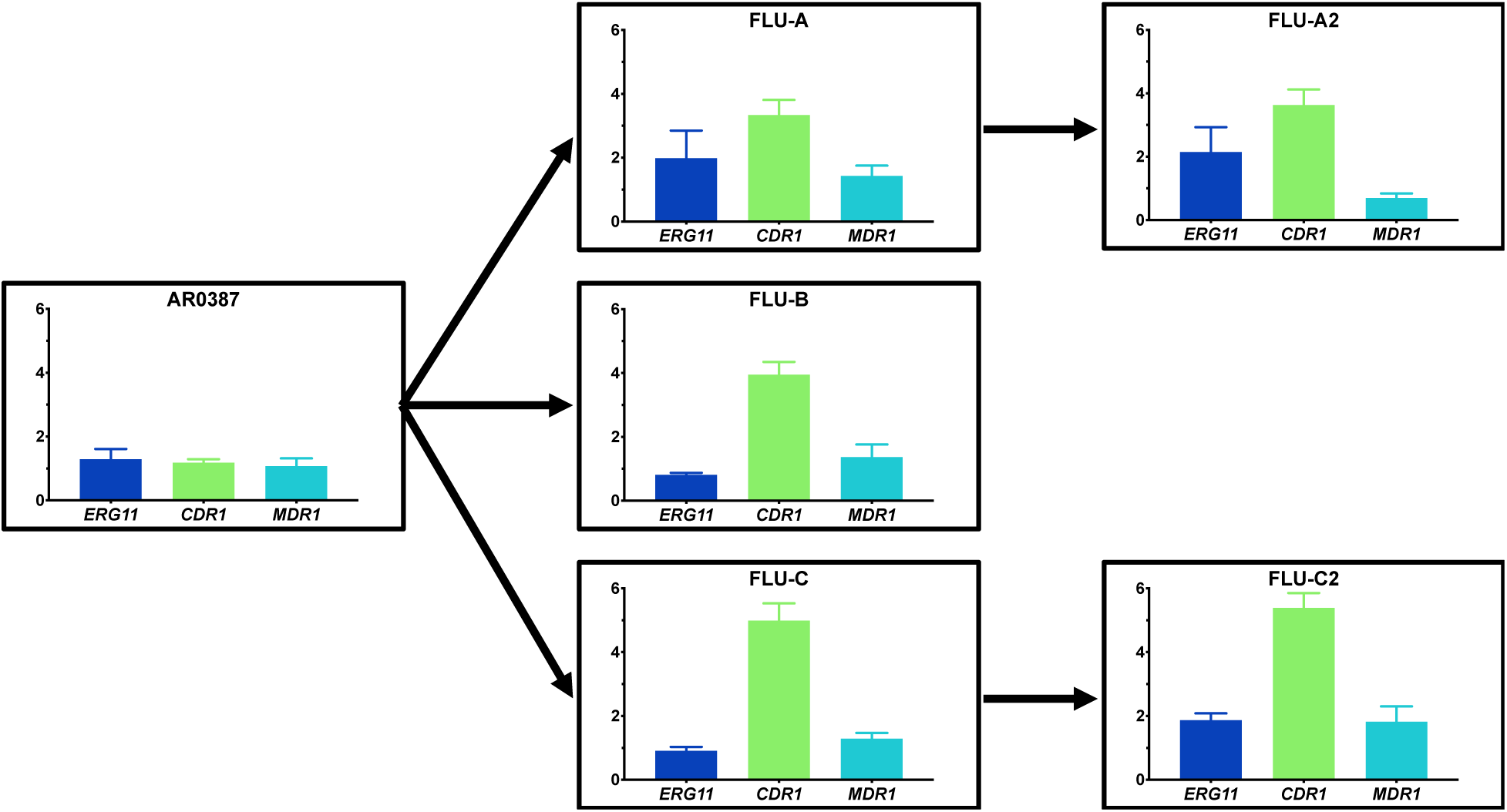
Relative expression of *C. auris ERG11, CDR1*, and *MDR1* among fluconazole-evolved strains grown to exponential growth phase in RPMI. The expression of *C. auris ERG11, CDR1*, and *MDR1* among AR0387 and the fluconazole-evolved strains were determined following culturing to exponential growth phase at 30°C in RPMI. The expression level for each sample is shown relative to that of the respective gene in AR0387. Arrows between graphs indicate lineage of each fluconazole-evolved strain from the parental AR0387.

As copy number variations (CNV) among genes encoding fluconazole resistance effectors, such as *ERG11*, have previously been reported among clinical isolates and laboratory strains of *C. auris*, qPCR amplifying from genomic DNA was performed to assess for CNV among the effectors *ERG11, CDR1*, and *MDR1*, as well as *TAC1B*, for each of the fluconazole-evolved strains. (26, 27) For each gene of interest, three primer sets spanning the open reading frame were utilized. While no alteration in the copy number of *ERG11, CDR1*, or *MDR1* was observed, the second-generation fluconazole-evolved strain FLU-A2 was found to have a 2-fold increase in the copy number of *TAC1B*, which was not evident in other evolved strains (**Figure S1**).

### *TAC1B* mutations identified during *in-vitro* evolution studies are also present among fluconazole-resistant *C. auris* clinical isolates

Interrogation of a dataset consisting of whole genome sequencing data for 304 globally distributed *C. auris* isolates representing each of the four major clades revealed mutations in *TAC1B* to be present among 164 isolates (54%).(27) Excluding sites which are fixed in all isolates within a clade, which are present in both sensitive and resistant isolates, 14 non-synonymous *TAC1B* mutations and one deletion, were identified (**Figure 8, Table S1**). Furthermore, the two *TAC1B* mutations that arose during *in-vitro* drug selection were found to be present among fluconazole-resistant clinical *C. auris* isolates, suggesting a possible role in clinical fluconazole resistance. R495G was found in a single Clade I isolate, and the F214S change was found in 2 isolates from Clade II and 1 isolate from Clade IV (Figure 8, Table S1). Notably, a mutation encoding the A640V amino acid substitution was found to be the most common among clinical isolates, found in 57 Clade I isolates from 7 countries and always present with the *ERG11* mutation encoding the K143R amino acid substitution. Nearly all (98.2%) of the isolates isolates with A640V and K143R mutations displayed high-level fluconazole resistance (>64mg/L). Other common *TAC1B* mutations include A657V in 15 Clade I isolates and a frame shift mutation F862_N866del in 46 Clade IV isolates. These mutations appeared in isolates with the *ERG11* Y132F variant, and these isolates have markedly high MIC values (Figure 8), suggesting these mutations may provide additive fluconazole resistance. Comparison of Tac1B protein sequences indicated *C. auris* A657V corresponds to the CaTac1 GOF mutation A736V which is associated with increased triazole resistance in *C. albicans*. Additionally, we observed three novel *TAC1B* mutations in Clade IV isolates lacking resistance-associated mutations in *ERG11*, including K247E (n = 5), M653V (n = 7) and A651T (n = 16), six resistant isolates from Clade I which harbored two *TAC1B* mutations (A15T and S195C), as well as two different mutations affecting the P595 site (P595L in Clade I and P595H in Clade IV).

**Figure 8.**
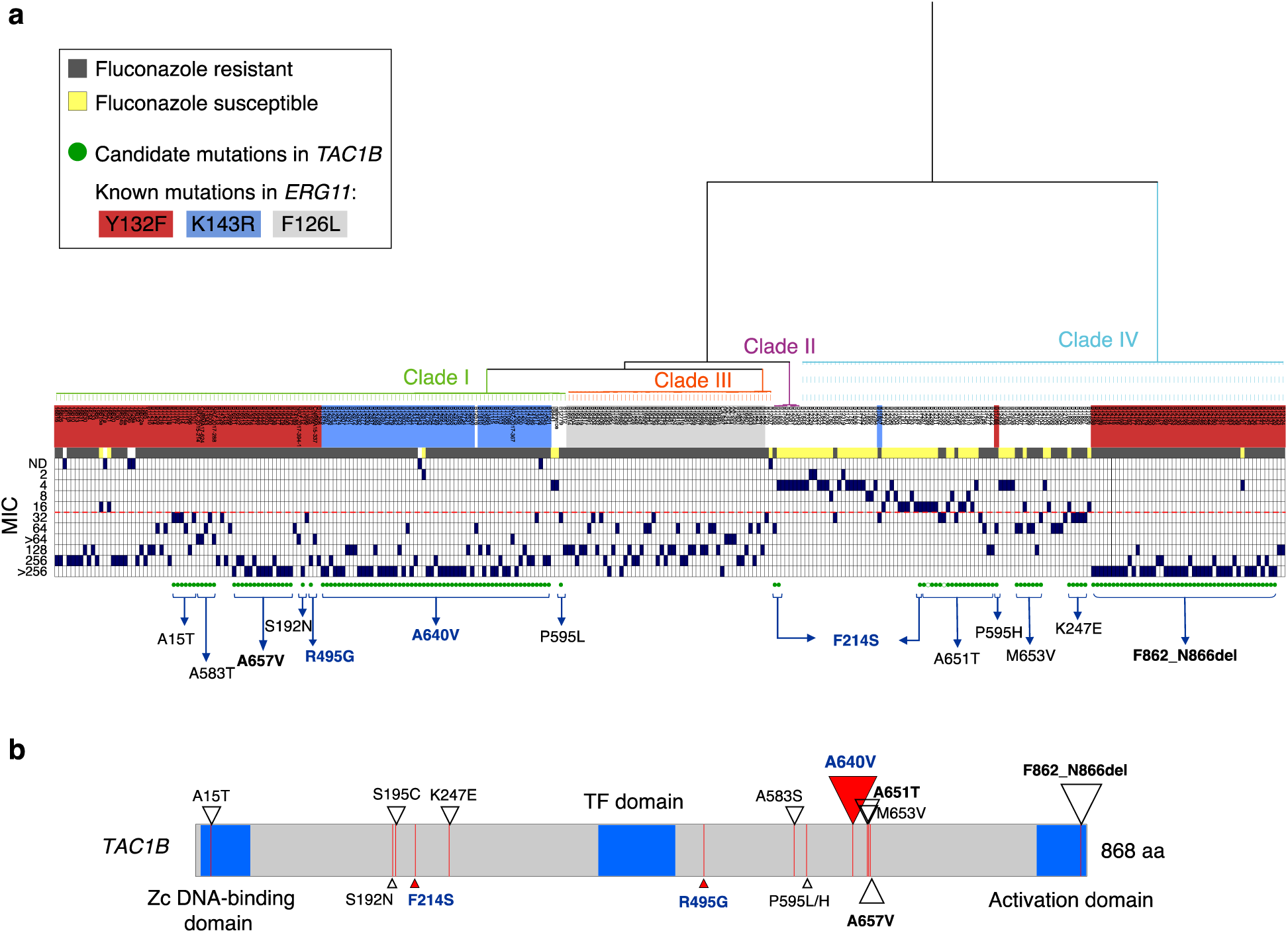
Point mutations *TAC1B* and fluconazole susceptibility in *C. auris*. (a) Phylogenetic tree of SNPs identified from 304 C. auris whole-genome sequences from four major clades (I, II, III and IV). Isolate label background are color-coded by known mutations in *ERG11* (B9J08_001448)(Y132F, K143R, F126L). Susceptibility to fluconazole is depicted as resistant (dark-gray) or susceptible (yellow), and the minimal inhibitory concentration (MIC) value is indicated as dark-blue boxes. The red dotted line indicates the tentative MIC breakpoint to fluconazole (≥32 mg/L). Green circles indicate isolates harboring non-clade specific non-synonymous mutations or gain-of-function mutations in *TAC1B* (B9J08_004820), with filled circles corresponding to percent alternate allele of ≥0.8 while open green circles correspond to percent alternate allele of 0.67 to 0.79. The specific mutation is indicated for each isolate(s).Mutations in bold/dark-blue arose in *in-vitro* evolution experiments or were functionally tested in this study and associated with increased resistant to fluconazole in *C. auris*. (b) Mutations and location in *TAC1B* protein sequence associated with azole resistance are indicated using triangles. Mutations in bold/dark blue (red triangles) arose in *in-vitro* evolution experiments or were functionally tested in this study and associated with increased resistant to fluconazole in *C. auris*. The size of the triangle indicates the number of isolates from this study harboring the mutation (range: 1 to 57 isolates).

### Mutations in *TAC1B* contribute to fluconazole resistance

As mutations in *TAC1B* were identified among a large proportion of fluconazole-resistant *C. auris* clinical isolates, and the mutation encoding the amino acid substitution A640V was found to be the most prevalent among this large collection of clinical isolates, the direct impact of this mutation on fluconazole susceptibility was next determined using a Cas9-mediated transformation system. To accomplish this, the *TAC1B* allele from the previously characterized fluconazole-resistant *C. auris* clinical isolate AR0390 (also known as B11205, an isolate from Clade I), which contains the mutation encoding the amino acid substation A640V, was introduced into the fluconazole-susceptible clinical isolate AR0387 using the Cas9-ribonucleoproteins (Cas9-RNP) and the *SAT-FLP* system as previously described.(20) Two independent positive transformants were obtained, and fluconazole MIC were determined by broth microdilution. Introduction of the *TAC1B*^A640V^ allele to the native *TAC1B* locus was observed to increase fluconazole MIC 8-fold relative to the parental AR0387 (**Figure 9**). Conversely, when the same methods were used to introduce the wildtype *TAC1B* allele to isolate AR0390 (which harbors the *TAC1B* mutation encoding A640V), a 16-fold decrease in fluconazole MIC was observed (**Figure 9**). Fluconazole MIC did not differ between independent transformants.

**Figure 9.**
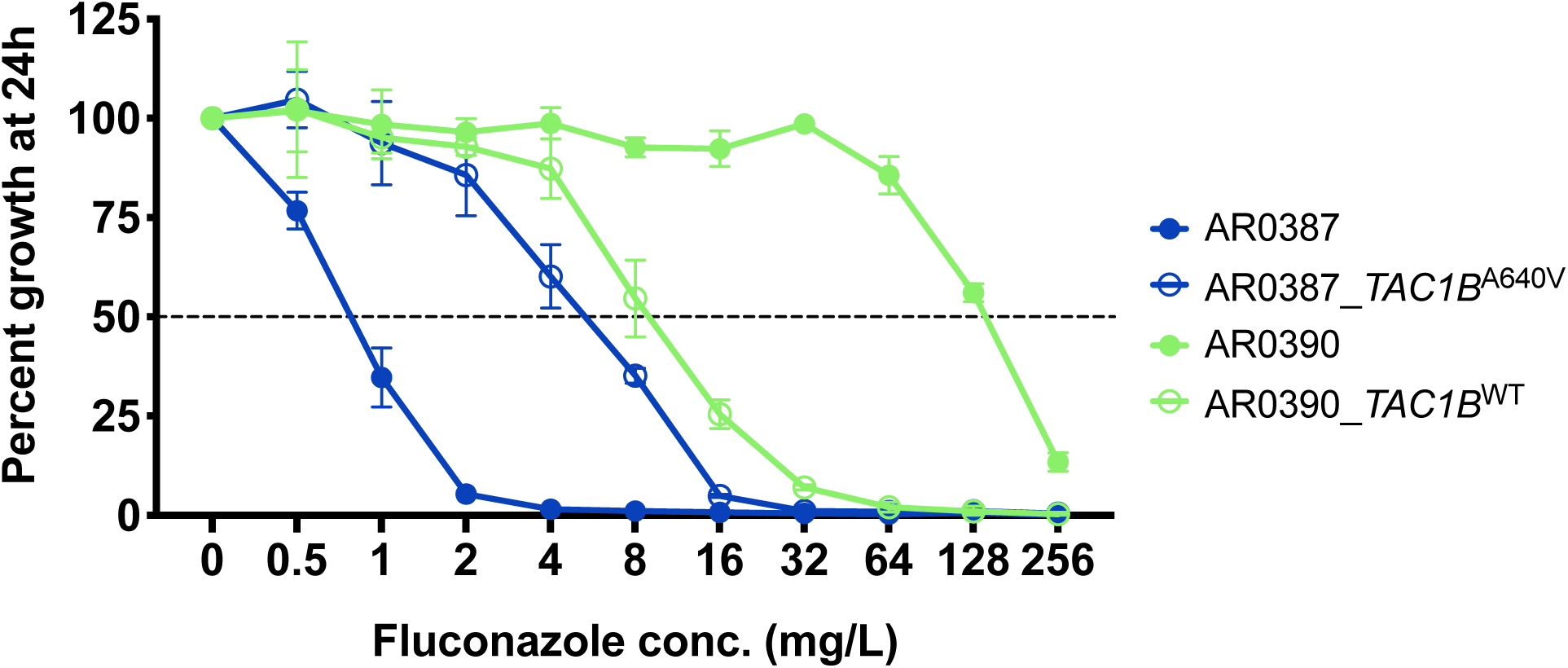
Fluconazole MIC for *TAC1B* strains. Percentage growth of AR0387, AR0390, and their derivative *TAC1B* strains with escalating concentrations of fluconazole measured at 24 hours. Percent growth was determined relative to respective untreated controls as assessed by absorbance at OD_600_.

## DISCUSSION

*C. auris* has rapidly become a fungal pathogen of global concern. Among the characteristics most notably distinguishing this organism from other species of *Candida*, the prevalence of fluconazole resistance is of clear clinical concern. While mutations in the *ERG11* gene are strongly associated with clinical fluconazole resistance in *C. auris*, other genetic and molecular mechanisms contributing to fluconazole resistance in this organism are largely unknown.

In this work, we utilize both *in vitro* evolution and large-scale whole genome sequencing to identify mutations in *TAC1B* as a novel genetic determinant of fluconazole resistance. Mutations in *TAC1B* were identified among all fluconazole-evolved strains, and associated with altered sterol profiles, decreased [^3^H]-fluconazole uptake, and increased expression of *CDR1* and *TAC1B*. A large-scale analysis of whole genome sequencing data for over 300 *C. auris* clinical isolates revealed the majority of isolates to harbor mutations in *TAC1B*, including those mutations identified among fluconazole-evolved strains as well as mutations similar to known GOF mutations in *C. albicans TAC1*. Subsequently, Cas9-RNP mediated genetic manipulations demonstrated that the most common mutation found among fluconazole-resistant clinical isolates of *C. auris*, encoding the amino acid substitution A640V, alone was sufficient to elevate fluconazole resistance by 8-fold. Thus, mutations in *TAC1B* are both a potent genetic determinant contributing to clinical fluconazole resistance in *C. auris*, and prevalent among a large global collection of fluconazole-resistant clinical isolates. Further studies characterizing the interplay between mutations in *ERG11* and *TAC1B*, and the delineation of the *TAC1B* regulon in *C. auris* are needed.

## METHODS

### Isolate, strains, and growth media used in this study

Clinical isolates AR0387 and AR0390 were made available by the CDC and FDA AR Isolate Bank as part of the *C. auris* collection of isolates. All constructed strains and clinical isolates were grown in YPD liquid media (1% yeast extract, 2% peptone, and 2% dextrose) at 30°C in a shaking incubator unless otherwise indicated. Frozen stocks of all strains and clinical isolates were prepared with 50% sterile glycerol and were maintained at −80°C.

### Minimum inhibitory concentration determination

Fluconazole (Sigma) was prepared in DMSO. As previously described, modified Clinical and Laboratory Standards Institute document M27 methodology utilizing broth microdilution, RPMI liquid media, and reading absorbance at 600 nm on a BioTek Synergy 2 microplate reader (BioTek, Winooski, VT), were used to determine fluconazole MIC as the lowest concentration at which 50% inhibition of growth was obtained.(28) All susceptibility testing was performed technical triplicate and biological duplicate.

### Comprehensive sterol profiling

Fluconazole-evolved strains and the parental clinical isolate were grown to exponential growth phase at 30°C in RPMI liquid media with or without fluconazole supplemented at 16mg/L. Alcoholic KOH was used to extract nonsaponifiable lipids. A vacuum centrifuge (Heto) was used to dry samples, which were then derivatized by adding 100 µl 90% N,O-bis(trimethylsilyl)-trifluoroacetamide–10% tetramethylsilane (TMS) (Sigma) and 200 µl anhydrous pyridine (Sigma) while heating at 80°C for 2 hours as previously described.(23, 29) GC-MS (with a Thermo 1300 GC coupled to a Thermo ISQ mass spectrometer; Thermo Scientific) was used to analyze and identify TMS-derivatized sterols through comparison of the retention times and fragmentation spectra for known standards. Sterol profiles for each sample were determined by analyzing the integrated peak areas from GC-MS data files using Xcalibur software (Thermo Scientific). All sterol analysis was performed in biological triplicate.

### Assessment of [^3^H]-fluconazole uptake

*C. auris* isolates and fluconazole-evolved strains were glucose-starved for 3 hours, and 200 µL of concentrated cell pellets were added to 250 µL of YNB without glucose and 50 µL of freshly diluted 0.77 µM [^3^H]-fluconazole, yielding a final [^3^H]-fluconazole concentration significantly below the MIC of each strain or isolate being tested (23.6 pg/L). Samples were then incubated at 30°C for 24 hours before 200 µL of each sample was transferred to 5mL of stop solution (YNB +20 mM [6mg/L] unlabeled fluconazole) in a 14mL round bottom tube. Samples were then filtered and dried on glass fiber filters, then washed with another 5mL of stop solution before the filters and cells were then transferred to a 5mL scintillation vial. A Beckman Coulter scintillation analyzer was then used to quantify the radioactivity of each filter following the addition of 3 ml of scintillation cocktail (Ecoscint XR, National Diagnostics). Experiments were performed with six biological replicates and all results were normalized to CPM per 1×10^8^ cells.

### Assessment of copy number variation by qPCR and relative gene expression by reverse transcription quantitative PCR

For assessment of gene copy number variation, genomic DNA was isolated from each isolate or strain, and qPCR was performed three independent primer sets spanning the open reading frame of each gene of interest, and the housekeeping gene *ACT1*, using SYBR green per the manufacturer’s instructions and as previously described.(26) For assessment of relative gene expression, *C. auris* isolates and strains were inoculated into 2 mL of RPMI broth buffered with MOPS to pH 7.0 and grown overnight at 30°C for initiation. Overnight cultures were then diluted to an OD_600_ of 0.1 in 10 mL of RPMI media with or without 16 mg/mL of fluconazole and placed in a 50 mL conical tube. Cultures were then incubated for 10 hours, confirmed to be exponential growth phase under these conditions, and then cells were collected by centrifugation, storing cell pellets at −80°C until isolation of RNA was performed. Synthesis of cDNA was performed using the RevertAid RT kit (Thermo Scientific), per the manufacturer’s instructions. *C. auris ACT1, ERG11, CDR1*, and *MDR1* were then amplified from cDNA using SYBR green, PCR master mix, and parameters as previously described.(20) All experiments were performed in biological and technical triplicate. The 2^-ΔΔ*CT*^ method was used to calculate relative expression of each gene of interest, and standard error was determined using Δ*CT* values as previously described.(22, 30) Primers are listed in **Table S2**.

### Variant identification

*TAC1A* (B9J08_004819) and *TAC1B* (B9J08_004820) mutations were identified in a set of 304 globally distributed *Candida auris* isolates representing Clade I, II, III and IV(27). For this dataset read quality and filtering was performed using FastQC v0.11.5 and PRINSEQ v0.20.3 (21278185) using “-trim_left 15 -trim_qual_left 20 -trim_qual_right 20 -min_len 100 -min_qual_mean 25 -derep 14”. Then, paired-end reads were aligned to the *C. auris* assembly strain B8441 (GenBank accession PEKT00000000.2; (30559369)) using BWA mem v0.7.12 (19451168) and variants were identified using GATK v3.7 (20644199) with the haploid mode and GATK tools (RealignerTargetCreator, IndelRealigner, HaplotypeCaller for both SNPs and indels, CombineGVCFs, GenotypeGVCFs, GatherVCFs, SelectVariants, and Variant Filtration). Sites were filtered with Variant Filtration using “QD < 2.0 ‖ FS > 60.0 ‖ MQ < 40.0”. Genotypes were filtered if the minimum genotype quality < 50, percent alternate allele < 0.8, or depth < 10 (https://github.com/broadinstitute/broad-fungalgroup/blob/master/scripts/SNPs/filterGatkGenotypes.py). Genomic variants were annotated and the functional effect was predicted using SnpEff v4.3T (22728672). The annotated VCF file was used to determine the genotype of known mutation sites in *ERG11* (B9J08_001448) and mutations in *TAC1A* (B9J08_004819) and *TAC1B* (B9J08_004820).

### Antifungal susceptibility testing for global collection of isolates

Fluconazole susceptibility testing was included for 294 of 304 isolates included in whole genome analyses. A total of 270 isolates were tested at the CDC as outlined by Clinical and Laboratory Standards Institute guidelines. Briefly, custom prepared microdilution plates (Trek Diagnostics, Oakwood Village, OH, USA) were used for fluconazole. Resistance to fluconazole was set at ≥32 mg/L. This interpretive breakpoint was defined based on a combination of these breakpoints and those established for other closely related *Candida* species, epidemiologic cutoff values, and the biphasic distribution of MICs between the isolates with and without known mutations for antifungal resistance (https://www.cdc.gov/fungal/candida-auris/c-auris-antifungal.html).

### Cas9-Ribonucleoprotein mediated transformations

*C. auris* Cas9 and electroporation-mediated transformations were performed as previously described with minor modification.(20) The *C. auris TAC1B* alleles from AR0387 (*TAC1B*^WT^) and AR0390 (*TAC1B*^A640V^) were amplified from genomic DNA, then cloned into the plasmid pBSS2 using restriction enzymes SacII and EagI, yielding the plasmids pBSS2-*TAC1B*^WT^ and pBSS2-*TAC1B*^A640V^. Repair templates for each allele of interest were then amplified from each plasmid using primers that also introduced approximately 50 bases of homology targeting the *TAC1B* loci to the 3’ end of the repair templates. Primers are listed in **Table S2**. Electrocompetent *C. auris* cells were prepared as previously described. Approximately 4 µM of dual Cas9-RNP that target both the *TAC1B* allele and the sequence immediately downstream of the open reading frame, as well as 1 µg of repair template were mixed with cells prior to electroporation according to the *C. albicans* protocol on a GenePulsar Xcell (Bio-Rad).(24) Cells were then allowed to recover for 4 to 6 hours in YPD, incubating in a shaking incubator at 30°C. Transformants were then selected by plating recovered cells on YPD plates supplemented with 400 µg/mL of nourseothricin. Integration of the repair template at the targeted loci was then confirmed by PCR for all transformants. The *FLP* recombinase was then induced by growing positive transformants in YPM (1% yeast extract, 2% peptone, and 2% maltose) to mediate loss of the *SAT1-FLP* cassette. All final strains identified to have lost the *SAT1-FLP* cassette by replica plating as previously described, were then again confirmed by sequencing.(18)

### Data and resource availability

All Illumina sequence analyzed in this project is available in the NCBI SRA under BioProjects PRJNA328792, PRJNA470683, PRJNA493622. A set of isolates are available from the CDC and FDA Antimicrobial Resistance (AR) Isolate Bank (https://www.cdc.gov/drugresistance/resistance-bank/index.html).

## ACKNOWLEDGMENTS

The authors thank the CDC for providing the *C. auris* isolates used in this study as part of the CDC & FDA Antibiotic Resistance Isolate Bank program. This work was supported by NIH NIAID grant R01 A1058145 awarded to P.D.R. CAC and JFM were supported by NIAID award U19AI110818 to the Broad Institute. CAC is a CIFAR fellow in the Fungal Kingdom Program.

## DISCLAIMER

The findings and conclusions in this report are those of the author(s) and do not necessarily represent the official position of the Centers for Disease Control and Prevention.

---

**Figure S1.**
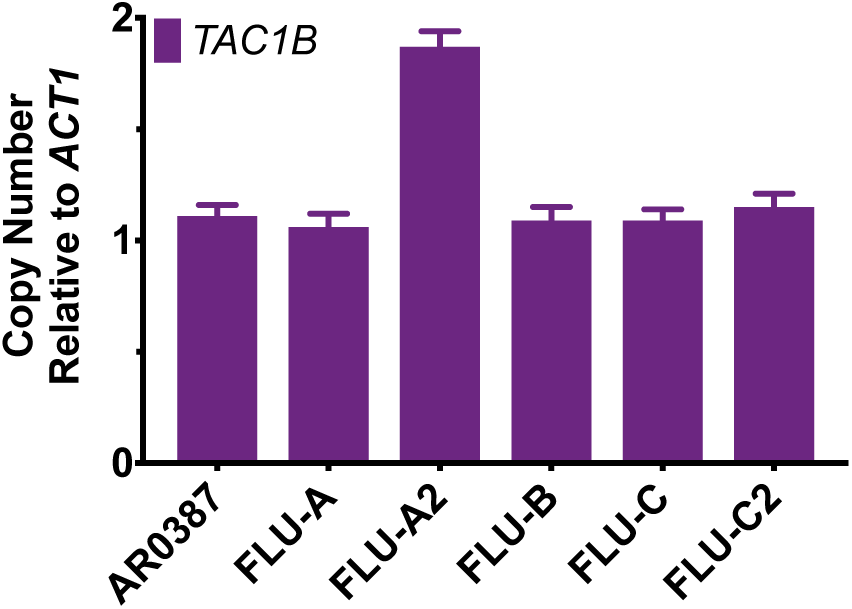
Copy number variation of *TAC1B* among fluconazole-evolved strains as determined by qPCR. Gene copy number of *TAC1B* across fluconazole-evolved strains and parental AR0387 as determined by qPCR with three independent primer sets spanning the open reading frame and as compared to the housekeeping gene *ACT1*.

## SUPPLEMENTARY TABLES

**Table S1.** *TAC1A* and *TAC1B* mutations observed among global collection of *C. auris* isolates. Variants shown in bold are unique to resistant isolates. Variants shown in orange were identified through *in vitro* evolution experiments. Variants shown in yellow were verified to directly contribute to fluconazole resistance. Variants shown in *italics* are uniquely found together. Percent shown represents percent of isolates with the indicated *TAC1B* mutation from the indicated Clade which have fluconazole MIC ≥32mg/L.

| GENE | VARIANT | Clade I<br>(n = 126)<br>%(n) | Clade II<br>(n = 7)<br>%(n) | Clade III<br>(n = 51)<br>%(n) | Clade IV<br>(n = 120)<br>%(n) | Total<br>n |
| --- | --- | --- | --- | --- | --- | --- |
| *TAC1B*<br>(B9J08_004820) | **A15T** | 100(6) | 0 | 0 | 0 | 6 |
|  | S36L | 0 | 0 | 51 | 120 | 171 |
|  | S89Y | 0 | 0 | 51 | 0 | 51 |
|  | S192N | 100(1) | 0 | 0 | 0 | 1 |
|  | **S195C** | 100(6) | 0 | 0 | 0 | 6 |
|  | E200K | 0 | 0 | 51 | 0 | 51 |
|  | **F214S** | 0 | 50(2) | 0 | 0(1) | 3 |
|  | K215R | 0 | 7 | 51 | 120 | 178 |
|  | K225N | 0 | 0 | 51 | 0 | 51 |
|  | Q226R | 0 | 0 | 51 | 120 | 171 |
|  | **K247E** | 0 | 0 | 0 | 80(5) | 5 |
|  | I268V | 0 | 0 | 51 | 0 | 51 |
|  | D278V | 0 | 0 | 51 | 120 | 171 |
|  | Q298K | 0 | 0 | 51 | 0 | 51 |
|  | L328Q | 0 | 0 | 0 | 120 | 120 |
|  | C331S | 0 | 0 | 51 | 120 | 171 |
|  | C334F | 0 | 0 | 51 | 120 | 171 |
|  | L335S | 0 | 0 | 51 | 120 | 171 |
|  | S339A | 0 | 0 | 51 | 120 | 171 |
|  | T346I | 0 | 0 | 51 | 0 | 51 |
|  | **R495G** | 100(1) | 0 | 0 | 0 | 1 |
|  | Q503R | 0 | 0 | 51 | 0 | 51 |
|  | F580L | 0 | 0 | 51 | 0 | 51 |
|  | **A583S** | 100(5) | 0 | 0 | 0 | 5 |
|  | **P595H** | 0 | 0 | 0 | 100(1) | 1 |
|  | **P595L** | 100(1) | 0 | 0 | 0 | 1 |
|  | P607S | 0 | 0 | 0 | 0(1) | 1 |
|  | Y608H | 0 | 0 | 51 | 120 | 171 |
|  | **A640V** | 98.2(57) | 0 | 0 | 0 | 57 |
|  | Y647C | 0 | 0 | 51 | 0 | 51 |
|  | A651T | 0 | 0 | 0 | 37.5(16) | 16 |
|  | M653V | 0 | 0 | 0 | 86(6) | 7 |
|  | **A657V** | 100(15) | 0 | 0 | 0 | 15 |
|  | S754N | 0 | 0 | 51 | 120 | 171 |
|  | M809I | 0 | 0 | 51 | 120 | 171 |
|  | N773_L774del | 0 | 0 | 51 | 0 | 51 |
|  | **F862_N866del** | 0 | 0 | 0 | 97.8(46) | 46 |
| TAC1A<br>(B9J08_004819) | V13I | 0 | 0 | 51 | 120 | 171 |
|  | S116A | 0 | 0 | 51 | 120 | 171 |
|  | V145E | 0 | 7 | 51 | 120 | 178 |
|  | G149D | 0 | 0 | 51 | 0 | 51 |
|  | A288S | 0 | 0 | 0 | 120 | 120 |
|  | E313G | 0 | 3 | 0 | 0 | 3 |
|  | P371L | 1 | 0 | 0 | 0 | 1 |
|  | D500E | 0 | 0 | 51 | 120 | 171 |
|  | E560D | 0 | 0 | 51 | 120 | 171 |
|  | E565D | 0 | 0 | 51 | 0 | 51 |
|  | S627G | 0 | 0 | 51 | 0 | 51 |
|  | K713N | 0 | 7 | 0 | 0 | 7 |
|  | E758G | 0 | 0 | 51 | 0 | 51 |
|  | S762P | 0 | 0 | 51 | 120 | 171 |
|  | A766T | 0 | 0 | 51 | 0 | 51 |

**Table S2.**
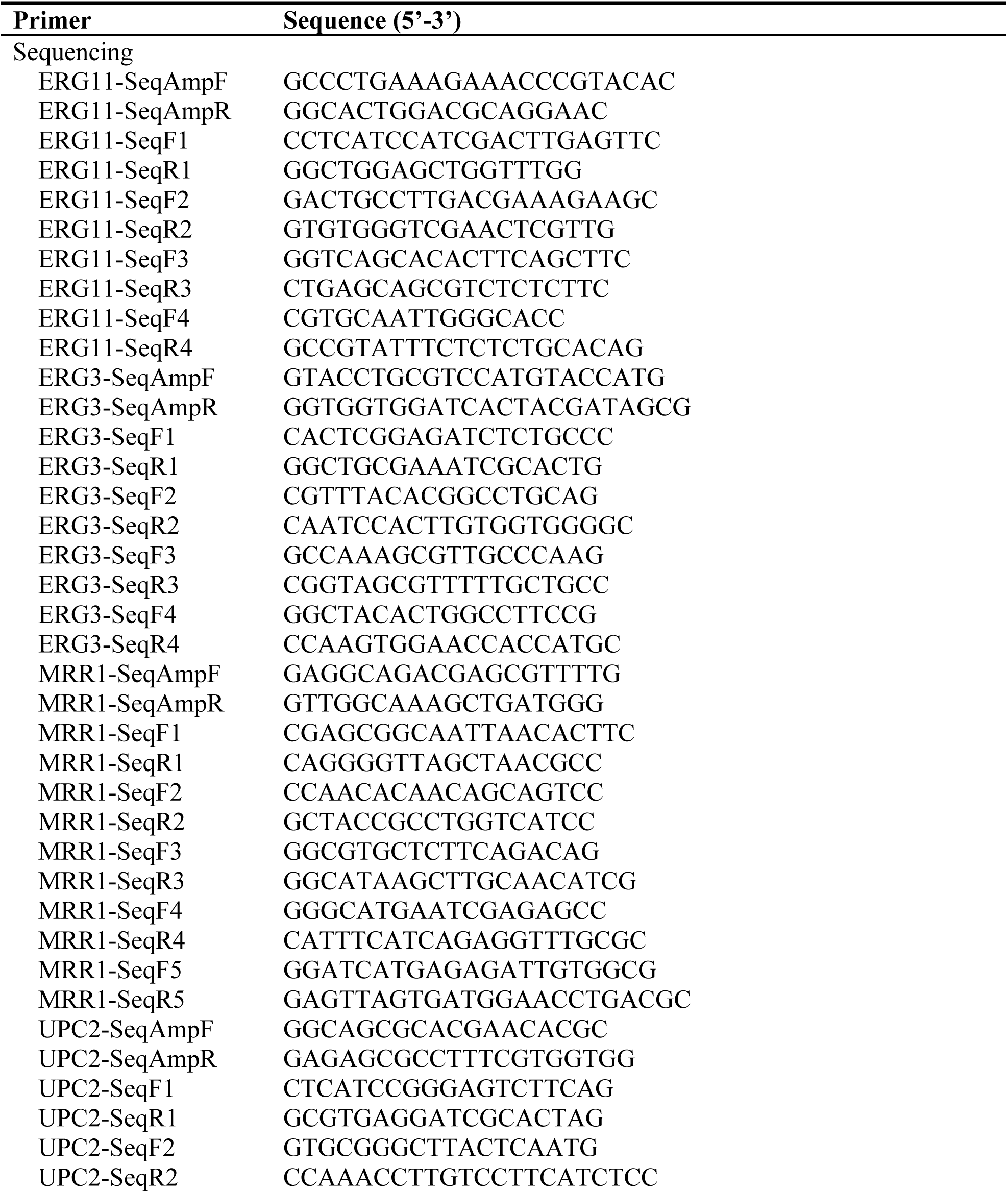
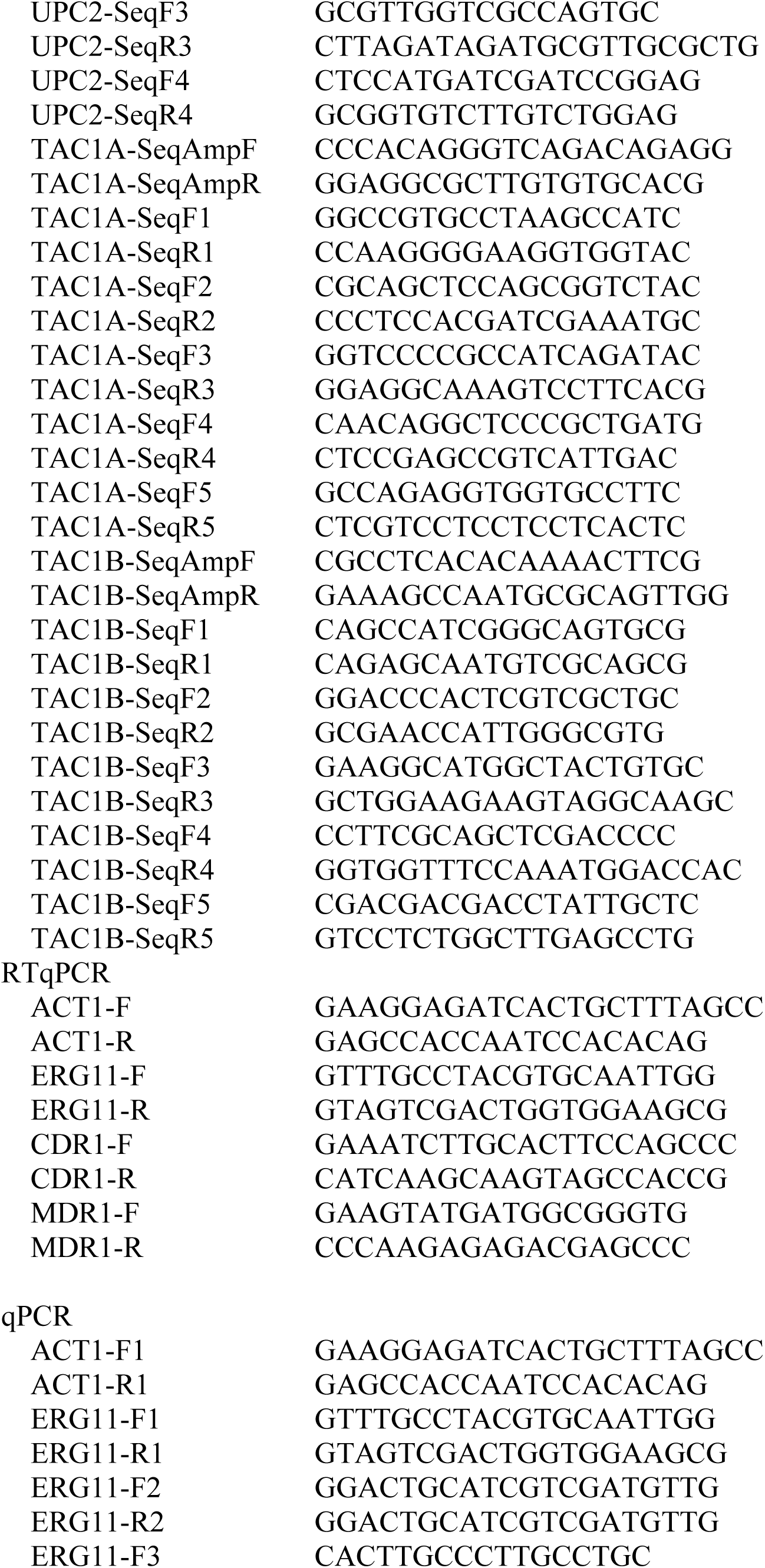
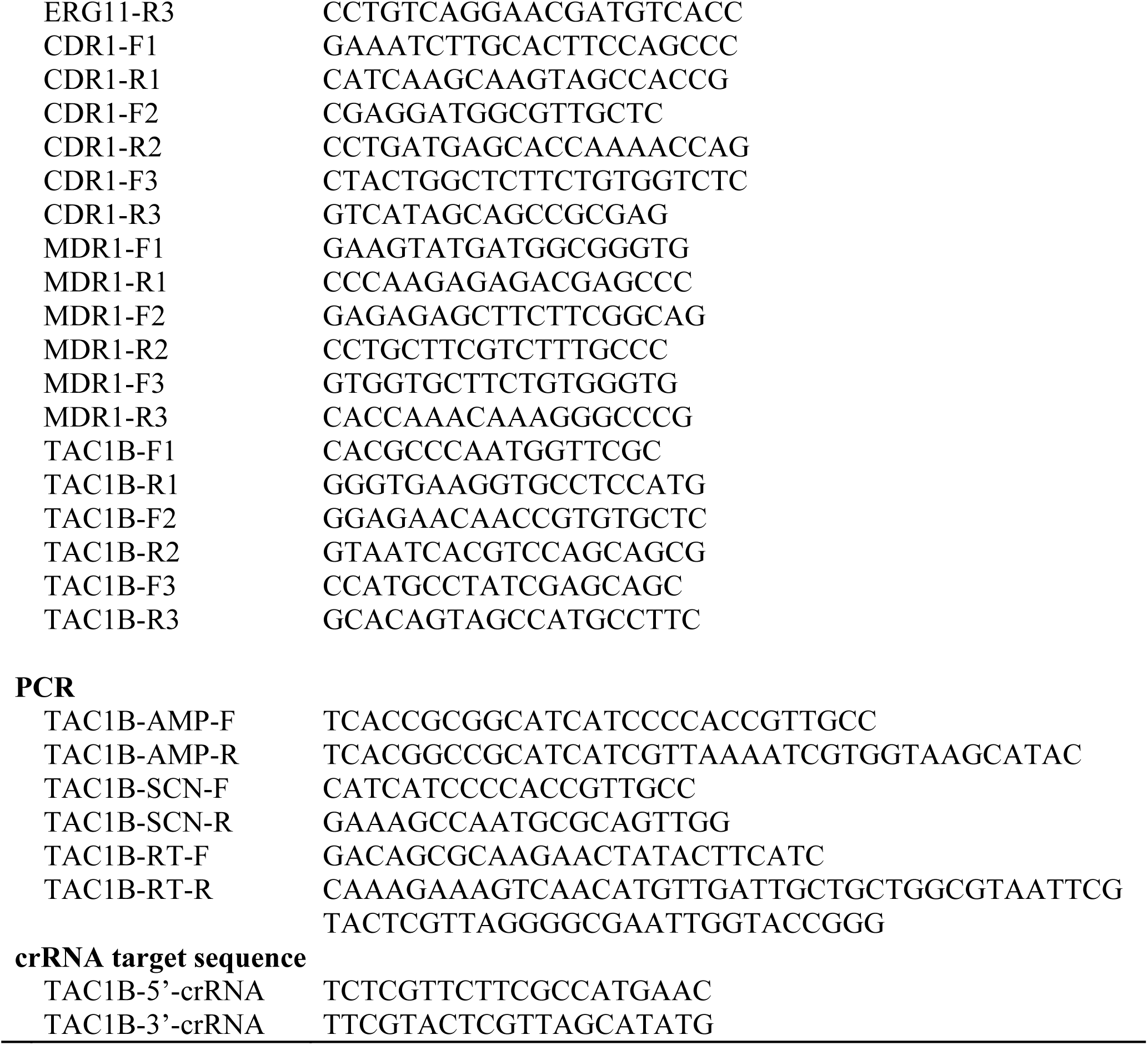
Oligonucleotides used in this study.

